# A molecular glue approach to control the half-life of CRISPR-based technologies

**DOI:** 10.1101/2023.03.12.531757

**Authors:** Vedagopuram Sreekanth, Max Jan, Kevin T. Zhao, Donghyun Lim, Jessie R. Davis, Marie McConkey, Veronica Kovalcik, Sam Barkal, Benjamin K. Law, James Fife, Ruilin Tian, Michael E. Vinyard, Basheer Becerra, Martin Kampmann, Richard I. Sherwood, Luca Pinello, David R. Liu, Benjamin L. Ebert, Amit Choudhary

**Affiliations:** Chemical Biology and Therapeutics Science Program, Broad Institute of MIT and Harvard, Cambridge, MA 02142, USA; Divisions of Renal Medicine and Engineering, Brigham and Women’s Hospital, Boston, MA 02115, USA; Department of Medicine, Harvard Medical School, Boston, MA 02115, USA; Broad Institute of Harvard and MIT, Cambridge, MA 02142, USA; Department of Medical Oncology, Dana-Farber Cancer Institute, Boston, MA 02215, USA; Department of Pathology, Massachusetts General Hospital, Boston, MA 02114, USA; Merkin Institute of Transformative Technologies in Healthcare, Broad Institute of Harvard and MIT, Cambridge, MA, USA; Department of Chemistry and Chemical Biology, Harvard University, Cambridge, MA, USA; Howard Hughes Medical Institute, Harvard University, Cambridge, MA, USA; Division of Genetics, Department of Medicine, Brigham and Women’s Hospital, Boston, MA 02115, USA; Institute for Neurodegenerative Diseases, Department of Biochemistry and Biophysics, University of California, San Francisco, San Francisco, CA 94158, USA; Chan-Zuckerberg Biohub, San Francisco, CA 94158, USA; Molecular Pathology Unit, Massachusetts General Hospital, Charlestown, MA, USA; Department of Pathology, Harvard Medical School, Boston, MA, USA; Broad Institute of Harvard and MIT, Cambridge, MA, USA; Hubrecht Institute for Developmental Biology and Stem Cell Research, Royal Netherlands Academy of Arts and Sciences (KNAW), Utrecht, The Netherlands; Howard Hughes Medical Institute, Dana-Farber Cancer Institute, Boston, MA 02215, USA

## Abstract

Cas9 is a programmable nuclease that has furnished transformative technologies, including base editors and transcription modulators (e.g., CRISPRi/a), but several applications of these technologies, including therapeutics, mandatorily require precision control of their half-life. For example, such control can help avert any potential immunological and adverse events in clinical trials. Current genome editing technologies to control the half-life of Cas9 are slow, have lower activity, involve fusion of large response elements (> 230 amino acids), utilize expensive controllers with poor pharmacological attributes, and cannot be implemented *in vivo* on several CRISPR-based technologies. We report a general platform for half-life control using the molecular glue, pomalidomide, that binds to a ubiquitin ligase complex and a response-element bearing CRISPR-based technology, thereby causing the latter’s rapid ubiquitination and degradation. Using pomalidomide, we were able to control the half-life of large CRISPR-based technologies (e.g., base editors, CRISPRi) and small anti-CRISPRs that inhibit such technologies, allowing us to build the first examples of on-switch for base editors. The ability to switch on, fine-tune and switch-off CRISPR-based technologies with pomalidomide allowed complete control over their activity, specificity, and genome editing outcome. Importantly, the miniature size of the response element and favorable pharmacological attributes of the drug pomalidomide allowed control of activity of base editor *in vivo* using AAV as the delivery vehicle. These studies provide methods and reagents to precisely control the dosage and half-life of CRISPR-based technologies, propelling their therapeutic development.

## Introduction

CRISPR-Cas9-based technologies, including those for base conversion (e.g., C→T using base editors) and transcription modulation (e.g., by using CRISPRi/a), are furnishing novel therapeutic modalities.^1–6^ Required of all therapeutic agents, the precision control of the half-life of CRISPR-based technologies is particularly crucial, as off-target editing, undesirable chromosomal translocations, genotoxicity, and activation of oncogenic pathways^7, 8^ are observed at the elevated or prolonged activities of these technologies^9–16^ and the longevity of Cas9 dictates genome editing outcome.^17, 18^ Finally, the pre-existence of immunity against Cas9 in humans will require its half-life control for therapeutic applications.^19, 20^

An ideal system to control the half-life of CRISPR-based technologies would have several characteristics. First, the system should be capable of degrading both small (< 10 kDa) and large (> 200 kDa) CRISPR-based technologies without impairing their activity. Second, the system should be minimalistic, ideally consisting of a controller that acts *via* a response element fused to CRISPR-based technology. Third, the response element should be miniature. For example, we recently reported a minimalistic system consisting of Cas9 fused to an FKBP variant (i.e., the response element), and a controller that triggers degradation of the fusion protein. However, this system is non-ideal because the FKBP fusions are large, adding ~230 amino acids, which not only reduces Cas9 activity but also significantly aggravate the viral packaging of Cas9 in Adeno-Associated Virus (AAV)-based delivery systems. Fourth, since contemporary and emergent CRISPR-based technologies already bear variable effector domains on Cas9 termini, a generalizable system should have a response element located internally on Cas9. Since our FKBP-system required termini fusion on Cas9, it did not afford control of CRISPR-based technologies, including CRISPRa/i and base editors. Fifth, the controller should be fast-acting to afford precision temporal control of the technology’s activity.^21^ Here, a small-molecule controller is preferable as they are fast-acting (vs. genetic methods)^22^ and the dose of the controller can be further used to control the half-life. However, the small molecule should be easily accessible and non-toxic; ideally, an FDA-approved drug will allow rapid translation of the system. Finally, since CRISPR-based technologies have diverse applications, the ideal system should be efficacious in diverse settings.

Leveraging rapid advancements in molecular glue technologies that induce targeted protein degradation, we report a general platform with the aforementioned characteristics that effectively controlled the half-life of both large and small CRISPR-based technologies. This platform utilizes a miniature response element (~60 amino acids) fused internally to Cas9 and the controller is pomalidomide, an FDA-approved drug.^23,24^ The response element and pomalidomide induce proximity between the CRISPR-based technology and ubiquitin ligase (Figure 1A), triggering the former’s ubiquitination and subsequent degradation by the host’s proteasome. This system can control the half-life of large technologies (e.g., Cas9, CRISPRi, base editors) and a small anti-CRISPR protein (~100 amino acids) that potently inhibits CRISPR-based technologies. By controlling the inhibitory activity of anti-CRISPRs using pomalidomide, we developed an “on-switch” for the base editor. Furthermore, pomalidomide allowed dose control of activity and/or specificity of Cas9, CRISPRi and base editors, as well as genome editing outcome. Finally, we demonstrated *in vivo* efficacy in mice and compatibility with AAV-delivery system using a base editor to knock-out *PCSK9,* a therapeutic target for hypercholesterolemia and cardiovascular diseases. Overall, we have developed a general and modular platform for precision control of half-life, activity, and specificity of CRISPR-based technologies in diverse cell types and *in vivo.*

**Figure 1.**
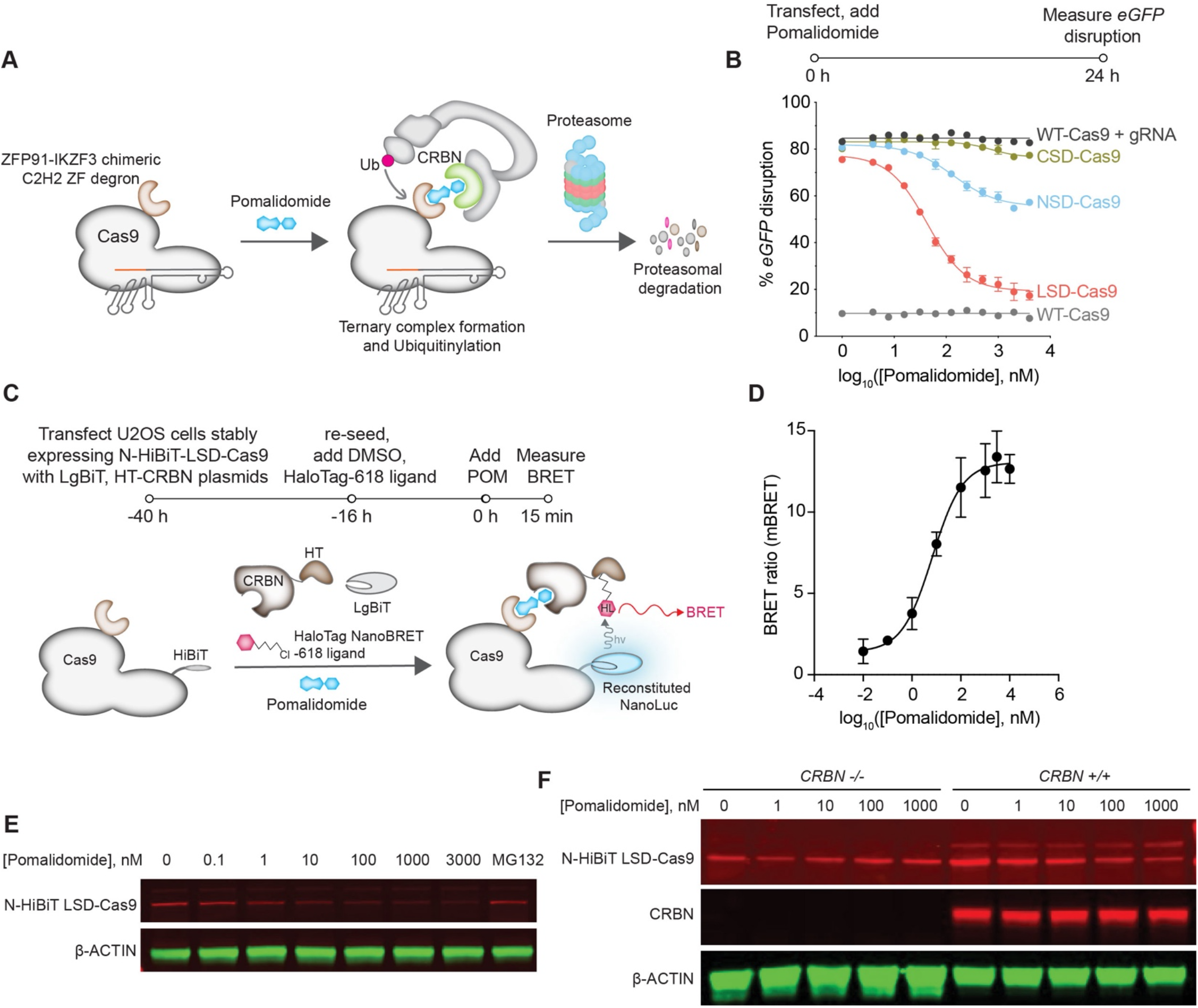
Demonstration of Cas9 degradation using superdegron derived from the short, 60-amino-acid pomalidomide-binding domain ZFP91-IKZF3. **(A)** Schematic showing the proteasomal degradation of Cas9 using the chimeric degron ZFP91-IKZF3 (superdegron) and pomalidomide. **(B)** Curves of pomalidomide dosedependent degradation of superdegron-Cas9 constructs in U2OS.eGFP.PEST cells in the *eGFP* disruption assay, made by analyzing by analyzing the images of the assay. **(C)** Schematic showing the NanoBRET based ternary complex formation between Halotag-cereblon (HT-CRBN), LgBiT reconstituted N-HiBiT-LSD-Cas9 and pomalidomide. **(D)** milliBRET ratio for the pomalidomide dose-induced ternary complex formation between LSD-Cas9 and HT-CRBN in N-HiBiT-LSD-Cas9 stably expressing U2OS cells. **(E)** Pomalidomide-induced degradation of N-HiBiT fused LSD-Cas9 in stably expressing U2OS cells. **(F)** Immunoblots showing the pomalidomide-induced degradation of N-HiBiT-fused LSD-Cas9 in transiently transfected HEK293T *CRBN -/-* and *CRBN +/+* cell lines.

## Results

### Control of half-life of Cas9

We fused Cas9 with a ~60-amino-acid, pomalidomide-binding response element called the superdegron (SD),^25,26^ at N-terminal, C-terminal, or loop-231 regions (NSD-, CSD-, and LSD-, respectively)^27, 28^ (Figure S1A). To test the activity of these fusion constructs, we used an eGFP-disruption assay wherein Cas9 knocks out *eGFP* and the concomitant loss of eGFP fluorescence reports on Cas9 activity.^29^ While all constructs showed a pomalidomide dose-dependent loss of Cas9 activity, the LSD-Cas9 had the largest dynamic range with complete loss of Cas9 activity at a pomalidomide concentration as low as 100 nM (Figure 1B, S1B).

Pomalidomide induces proximity between LSD-Cas9 and cereblon (CRBN), which triggers ubiquitination and degradation of the fusion protein. To demonstrate a ternary complex between LSD-Cas9: pomalidomide: and cereblon (CRBN), we used a bioluminescence resonance energy transfer (BRET) assay^30, 31^ wherein LSD-Cas9 bears a component of split luciferase (i.e., HiBiT) while the CRBN bears the other (i.e., LgBiT)—the ternary complex will reconstitute the luminescent activity of this split luciferase. The CRBN also bears a HaloTag (HT-CRBN) through which BRET-acceptor dye (i.e., NanoBRET-618 ligand) is conjugated (Figure 1C). U2OS cells stably expressing HiBiT-LSD-Cas9 were transfected with LgBiT-HT-CRBN and treated with NanoBRET-618 ligand. We observed a pomalidomide dose-dependent increased BRET signal, suggestive of ternary complex formation between HiBiT-LSD-Cas9, pomalidomide and LgBiT-HT-CRBN (Figure 1D). Since endogenous CRBN can also form the ternary complex and reduce signal, we implemented the BRET assay in *CRBN-/-* HEK293T cells and observed BRET signal even a 1 nM concentration of pomalidomide (Figure S1C).

To complement these assays, we performed immunoblotting in U2OS cells stably expressing N-HiBiT-LSD-Cas9 (Figure S1D) and observed significant degradation at concentrations as low as 100 nM (Figure 1E) and within 30 minutes of pomalidomide treatment (Figure S1E). To confirm that the degradation was CRBN mediated, we examined degradation of LSD-Cas9 in *CRBN* -/- and *CRBN* +/+ HEK293T cells and measured degradation using the split luciferase assay and immunoblotting. HEK293T cells were transfected with N-HiBiT-LSD-Cas9 plasmid and treated with different doses of pomalidomide for 24 h. The lysates were subjected to immunoblotting or treated with LgBiT followed by luminescence measurement.^32^ The immunoblotting experiments also confirmed CRBN mediated a dose-dependent decrease in the Cas9 levels in *CRBN* +/+ cells (Figure 1F). We observed no loss of luminescence signal from *CRBN* -/- cells, whereas *CRBN* +/+ cells yielded a pomalidomide dosedependent decrease in the luminescence signal (Figure S1F).

### Control of half-life of CRISPRi and base editors

Motivated by efficacious degradation of Cas9 by an internal response element, we applied this system to adenine base editors (ABEs) that can correct nearly half of known pathogenic point mutations by A•T to G•C conversion.^33^ We devised various constructs involving appending the response element to the deaminase, Cas9 nickase (nCas9) and the linker connecting the two (Figure S2A, S2B) and tested these constructs for A•T to G•C base conversion at *HBG2* gene.^34^ We find that the construct equivalent to LSD-Cas9 to be most efficacious while terminal other fusions [e.g., N-term to deaminase (ABE8e-SD1) or C-term fusion to nCas9 (ABE8e-SD7)] perturbed activity or were unresponsive to pomalidomide. Our optimal construct allowed significant degradation of the adenine base editor at as low as 100 nM of pomalidomide (Figure 2C, S2C), with most degradation being achieved in 8 h (Figure S2C, S2D).

**Figure 2.**
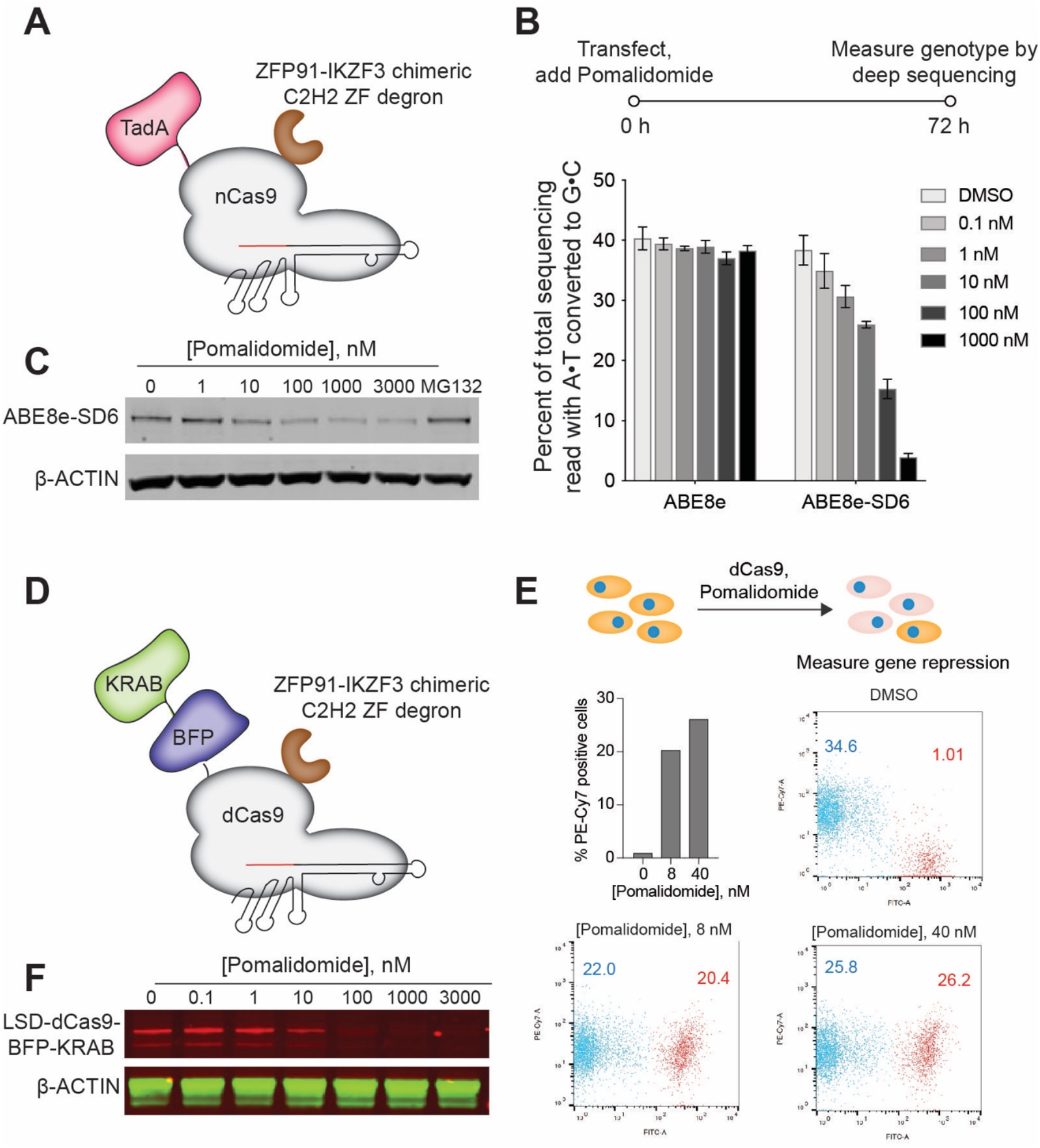
Generalizability of superdegron tags to base editors and CRISPRi systems. **(A)** Schematic of adenine base editor (ABE8e) fused with a single superdegron tag at loop-231 of the Cas9 nickase (ABE8e-SD6). **(B)** Pomalidomide dose-induced base-editor degradation in HEK293T cells transiently transfected with ABE8e and ABE8e-SD6 constructs. After 72 h of transfection and pomalidomide treatment, genomic DNA extracted was analyzed by NGS for the conversion of A·T to G·C. **(C)** Immunoblots showing pomalidomide-induced degradation of ABE8e-SD6 transiently transfected in HEK293T cells. **(D)** Schematic of LSD-dCas9-BFP-KRAB system. **(E)** A functional analysis of LSD-dCas9-BFP-KRAB upon pomalidomide-dependent degradation was carried out in iPSC cells by measuring the transferrin receptor (TFRC) protein levels via FACS. **(F)** Pomalidomide dose-induced dCas9 degradation in LSD-dCas9-BFP-KRAB stably expressing human iPSCs were monitored by immunoblotting.

Next, we implemented this degradation system to CRISPRi by appending the response element to a repressor construct (dCas9-BFP-KRAB, Figure 2D) at loop-231 of catalytically inactive dCas9 (i.e., equivalent site of Cas9 and base editors). To enable stable CRISPRi in induced pluripotent stem cells (iPSCs), we knocked in LSD-dCas9-BFP-KRAB into the citrate lyase beta-like (*CLYBL*) safe harbor locus, which enables robust transgene expression in iPSCs (Figure S2E).^35^ Once we established the stable iPSCs, we measured dose- and time-dependent degradation of dCas9 in the iPSCs via immunoblotting. Pomalidomide treatment at different concentrations yielded a dose-dependent degradation with the complete depletion at 100 nM of pomalidomide (Figure 2F) and a very short half-life of dCas9 of less than 30 minutes—the dCas9 fusion was completely degraded within 1 hour (Figure S2F).

We next appended the response element to a potent and strong repressor system, dCas9-KRAB-MeCP2. We transfected the HEK293T stable cells that are stably expressing WT- or LSD-dCas9-KRAB-MeCP2 cells (Figure S2G) with gRNAs targeting upstream of the C-X-C Motif Chemokine Receptor 4 (*CXCR4*) and Breast cancer type 1 (*BRCA1*) genes. We observed a dose-dependent increase in the expression of *CXCR4* and *BRCA1* genes, indicating the degradation of the dCas9-KRAB-MeCP2 system (Figure S2H).

We next tested the efficiency of dCas9/CRISPR gene silencing by evaluating the target protein levels in iPSCs stably expressing LSD-dCas9-BFP-KRAB. Because cell surface levels of transferrin can be easily and accurately quantified by antibody labeling and flow cytometry, we targeted the transferrin receptor (TFRC) gene, which encodes this ubiquitously expressed receptor responsible for iron uptake.^41^ We transfected the gRNA for TFRC upstream in the promoter region and observed severe depletion of the transferrin receptor protein. Pomalidomide treatment induced rapid degradation of LSD-dCas9-BFP-KRAB protein levels to restore the TFRC expression in iPSCs (Figure 2E). Overall, the highly controlled degradation of both base editors and CRISPRi using a simple plug-and-play system shows the transportability of our system across various CRISPR technologies in multiple cell lines, including iPSCs.

### Control of half-life of anti-CRISPRs to switch-on base editors

Controlled degradation of anti-CRISPR proteins that inhibit Cas9 can furnish a switch-on system for base editors allowing precise dose and temporal control of their activity. As AcrIIA4 is a potent anti-CRISPR, we fused the superdegron to the N- and C-terminus of AcrIIA4 (NSD- and CSD-) and tested their inhibitory activity in *eGFP* disruption assay, which pointed to CSD-AcrIIA4 as the best construct (Figure S3A, S3B). Use of a self-splicing linker connecting Cas9 to anti-CRISPR further enhanced the dynamic range and we observed a dose-dependent control of Cas9 activity by degrading the anti-CRISPR (Figure 3B). Using *HiBiT* knock-in assay in *CRBN* -/- and *CRBN* +/+ HEK293T cells, we showed that *HiBiT* luminescence was enhanced in a pomalidomide dose-dependent manner in *CRBN* +/+ cells but not in *CRBN* -/- cells, indicating the activation of Cas9 upon AcrIIA4 degradation (Figure S3C). We generated a CSD-AcrIIA4-P2A-eGFP stable cell line to precisely measure the degradation dose and kinetics (Figure S3D). Immunoblot analysis of dose and time-dependent degradation of CSD-AcrIIA4 in this line revealed that CSD-AcrIIA4 vanished at a pomalidomide concentration (D_max_) as low as 100 nM (Figure 3C, Figure S3E) and within ~25 minutes (DT_max_) (Figure 3D, Figure S3F).

**Figure 3.**
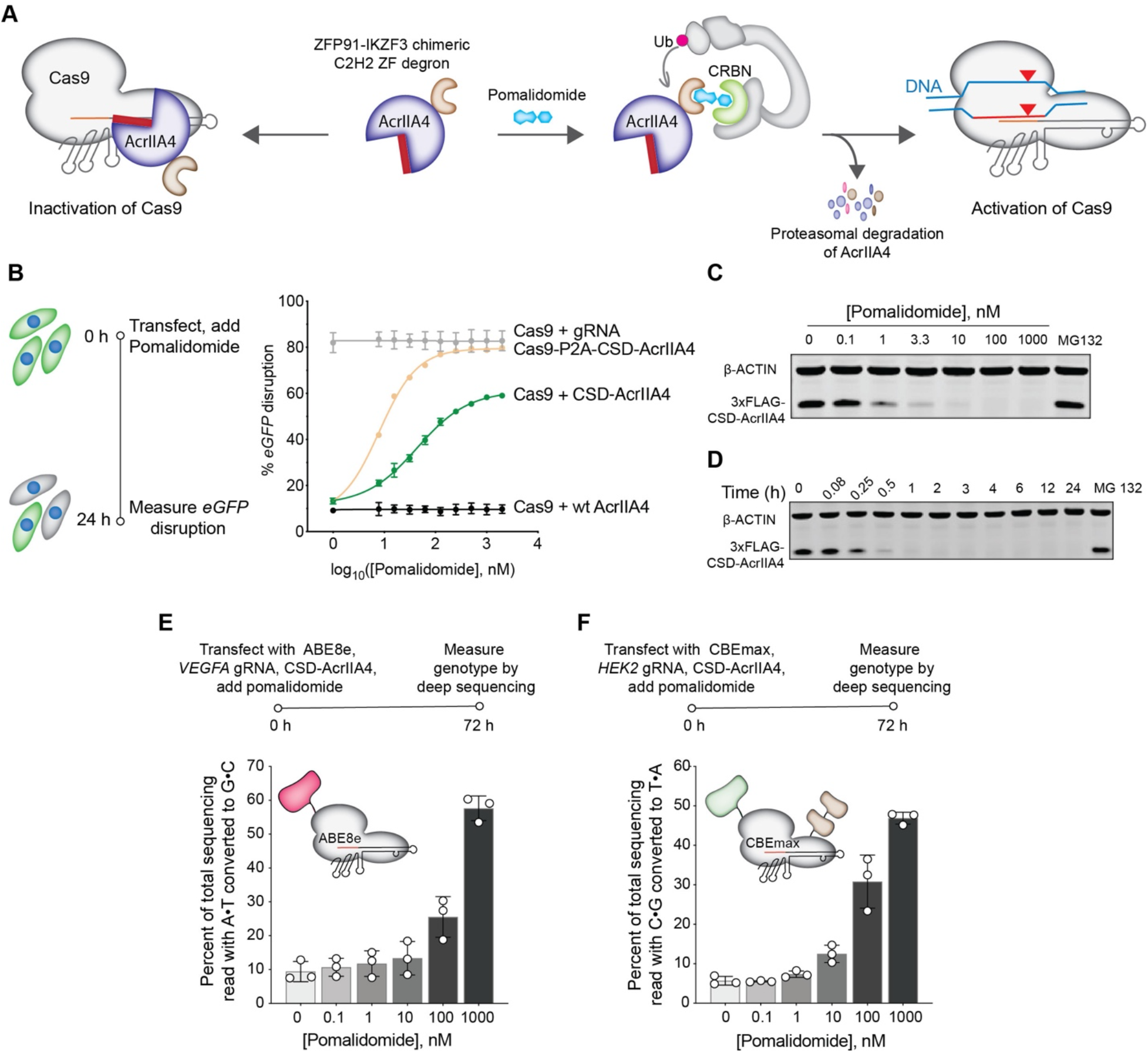
Demonstration of AcrIIA4 degradation-mediated Switch-On system for Cas9, base editor activation. **(A)** Schematic showing the proteasomal degradation of AcrIIA4 using the chimeric degron ZFP91-IKZF3 (superdegron) and pomalidomide leads to activation of Cas9. **(B)** The Cas9-P2A-CSD-AcrIIA4 fusion was investigated for pomalidomide-induced degradation using the *eGFP*-disruption assay. **(C, D)** Immunoblots for pomalidomide-induced dose- (C) and time-dependent (D) degradation of CSD-AcrIIIA4. **(E, F)** Pomalidomide dose dependent degradation of CSD-AcrIIA4 activates the adenine base editor (ABE8e) (E) and the cytosine base editor (CBEmax) (F) were measured by % conversion of A.T to G.C and % conversion of C.G to T.A base pair respectively by NGS.

Following the development of this robust and rapid anti-CRISPR degradation system, we endeavored to build an on-switch for adenine base editor (ABE8e) and cytosine base editor (CBEmax). Here, HEK293T cells were transfected with ABE8e (with *VEGFA* targeting gRNA) or CBEmax (with *HEK2* targeting gRNA) along with CSD-AcrIIA4 plasmids and treated with different concentrations of pomalidomide. We measured the editing efficiency A•T to G•C conversion (for ABE8e) or C•G to T•A (for CBE-max) conversion by next-generation sequencing. Both conversions indicate dose-dependent activation of adenine (Figure 3E) and cytosine base editor (Figure 3F) by pomalidomide. Overall, these studies offer a previously unavailable “on-switch” for base editors by controlling the levels of anti-CRISPR protein.

### Modulating DNA repair outcome by controlling Cas9 half-life

Following Cas9-induced double-strand break, the major repair pathways are non-homologous end joining (NHEJ) and microhomology-mediated end joining (MMEJ). NHEJ causes a smaller indel (1–4 nt deletions and insertions), while MMEJ causes deletions of variable sizes, so moderating the ratio between the repair pathways can offer control over the output genotype.^36^ To examine the effects of the half-life of Cas9 on the DNA repair outcomes, we assessed repair outcomes at a set of 48 target sites with corresponding paired gRNA that were genomically integrated into U2OS cells; this targetsite library is referred to as “Reduced Library”.^37^ The U2OS cells stably expressing the Reduced Library were transfected with a plasmid encoding LSD-Cas9 and following blasticidin selection, the cells were treated with pomalidomide at 6, 12, 24, and 48 hours and genomic DNA was collected at 120 hours. Non-MH deletions arising from the NHEJ pathway was favored over MH deletions arising presumably from the MMEJ pathway (Figure 4A). These results are consistent with the idea that the NHEJ pathway responds quickly to Cas9-mediated double-strand break and MH-deletion products accumulate with a prolonged Cas9 half-life.^27^ In addition to the changes in the relative frequency of MH deletion outcomes versus non-MH deletion outcomes, the observed frequency of 1 bp insertion products increased with prolonged LSD-Cas9 exposure (Figure 4A). Overall, these studies confirm that our system allows modulation of repair product outcomes by restricting the half-life of Cas9 in cells.

**Figure 4.**
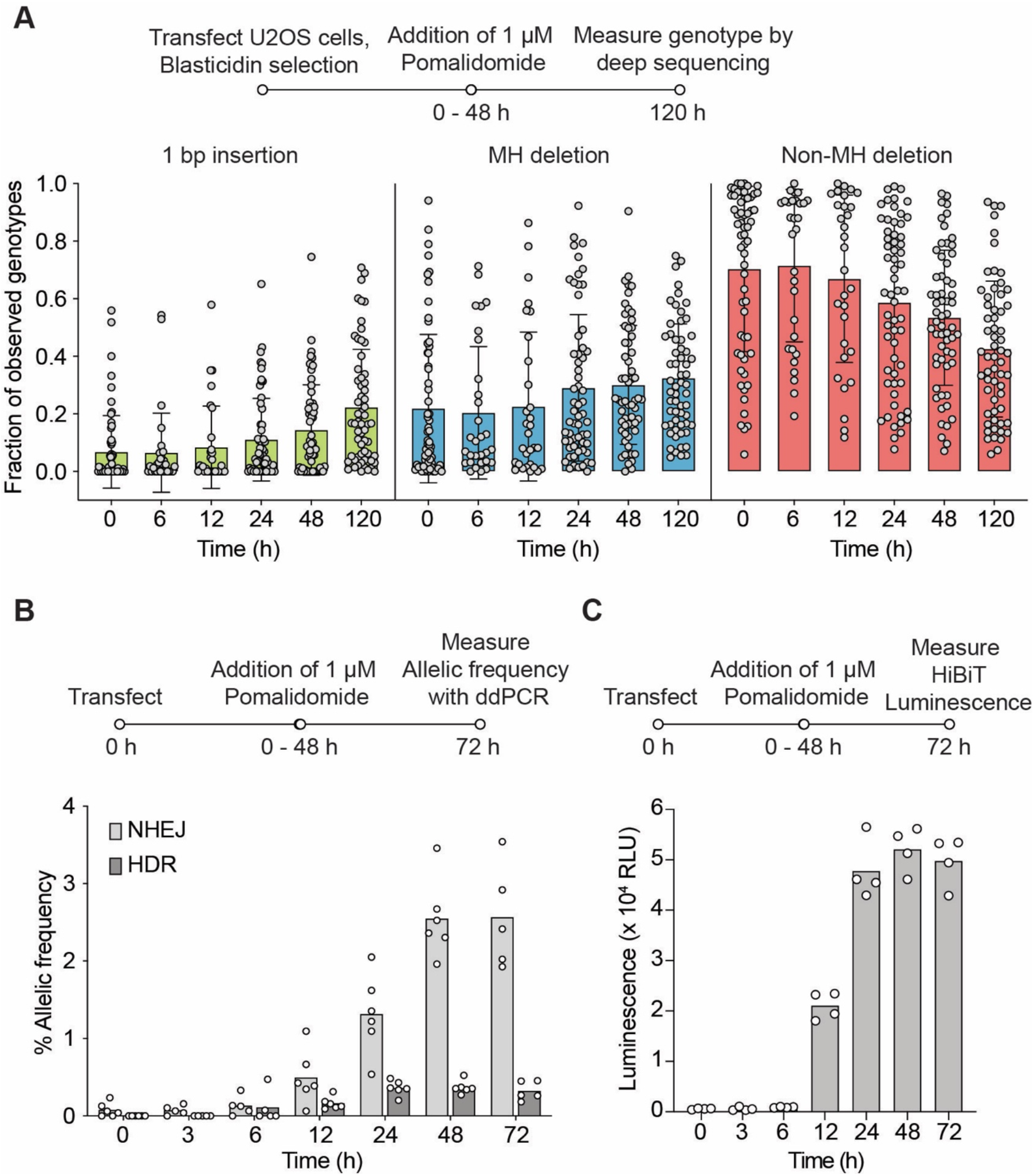
Cas9 half-life can impact DNA repair outcome. **(A)** U2OS cell line stably expressing the Reduced Library of 48 target sites used to test editing repair outcomes was transfected with the LSD-Cas9 plasmid and treated with 1 μM pomalidomide at different time points after transfection (0-48 h). The genomic DNA was extracted at 120 h post-transfection, and HTS sequencing was performed to analyze the +1 bp insertions, MH deletions, and non-MH deletions. **(B)** ddPCR quantification of single-nucleotide exchange at the *RBM20* locus in HEK293T cells following templated DNA repair. For this, LSD-Cas9 plasmid, *RBM20* gRNA plasmid, and ssODN template were transfected in HEK293T cells and were treated with pomalidomide at different time points after transfection. Cells were harvested at 72 h post-transfection, and percentages of HDR and NHEJ in the genomic DNA were analyzed by ddPCR analysis. (C) Luminescence-based quantification of *HiBiT* knock-in at the *GAPDH* locus in HEK293T cells following templated DNA repair. LSD-Cas9 plasmid, *GAPDH* gRNA plasmid, and ssODN template were transfected in HEK293T cells and were treated with pomalidomide at different time points after transfection. Cells were lysed at 72 h post-transfection and complemented with LgBiT protein to measure the luminescence.

Next, we evaluated the impact of modulation of the half-life of Cas9 on homology-directed repair (HDR), which is used for “knock-in” using an exogenously supplied donor DNA. Though a double-strand break can be repaired via a precise repair pathway (i.e., HDR), these products were much less frequent than the error-prone NHEJ- or MMEJ. Various approaches have been reported to enhance HDR frequency, including using small-molecule inhibitors of NHEJ,^38, 39^ though such approaches can have severe adverse effects.^40^ Here, we reasoned that modulating the half-life of Cas9 instead would provide better control over the relative levels of HDR and error-prone outcomes. To test this, we transfected HEK293T cells with plasmids encoding the LSD-Cas9 and RNA Binding Motif Protein 20 (*RBM20*) targeting gRNA along with a single-stranded oligonucleotide donor (ssODN).^27^ Pomalidomide was added at the indicated time points (0, 3, 6, 12, 24, and 48 h post-transfection), and the HDR and error-prone repair frequencies were investigated using droplet digital PCR (ddPCR).^41^ We observed the error-prone repair frequency increased with an increasing Cas9 half-life, while the HDR frequency saturated at about 24 h (Figure 4B).

We further evaluated the modulation of Cas9 half-life in the levels of HDR after the knock-in of a long ssODN.^27^ Here, HEK293T cells transfected with the LSD-Cas9 plasmid, Glyceraldehyde-3-Phosphate Dehydrogenase (*GAPDH*) targeting gRNA plasmid, and the HiBiT ssODN were treated with pomalidomide at the indicated time points (0, 3, 6, 12, 24, and 48 h post-transfection). At 72 hours post-transfection, cell lysates were complemented with LgBiT, and the HDR activity was quantified by HiBiT luminescence. Similar to the ddPCR results, the HiBiT HDR frequency saturated at about 24 h (Figure 4C). Our results suggest that a shortened Cas9 half-life or Cas9 inhibition offers a higher relative amount of “knock-in” product.

### Shortened Cas9 half-life enhances on-target specificity

As the off-target activity of Cas9 often displays slower kinetics than on-target activity, we reasoned that shortening Cas9 half-life should enhance specificity.^40, 41^ We measured the on- versus the off-target ratio of for Cas9. For each construct, cells were transfected, treated with pomalidomide at various doses or at different times, followed by extraction and sequencing of specific genomic sites to determine the on- versus off-target editing. For LSD-Cas9, we observed a dose-dependent reduction of off-target editing at the *EMX1* and *VEGFA* loci in HEK293T cells (Figure 5A, 5C and Figure S4A, S4C). We anticipated that a fast degradation of Cas9 will allow us to titer the optimal Cas9 half-life to maximize the on-target editing. At different time points after transfection with LSD-Cas9 and *EMX1-* or *VEGFA*-targeting gRNA, pomalidomide was added to HEK293T cells. We observed an enhanced on- versus off-target ratio with a shortened Cas9 half-life (Figure 5B, 5D and Figure S4B, S4D).^27^

**Figure 5.**
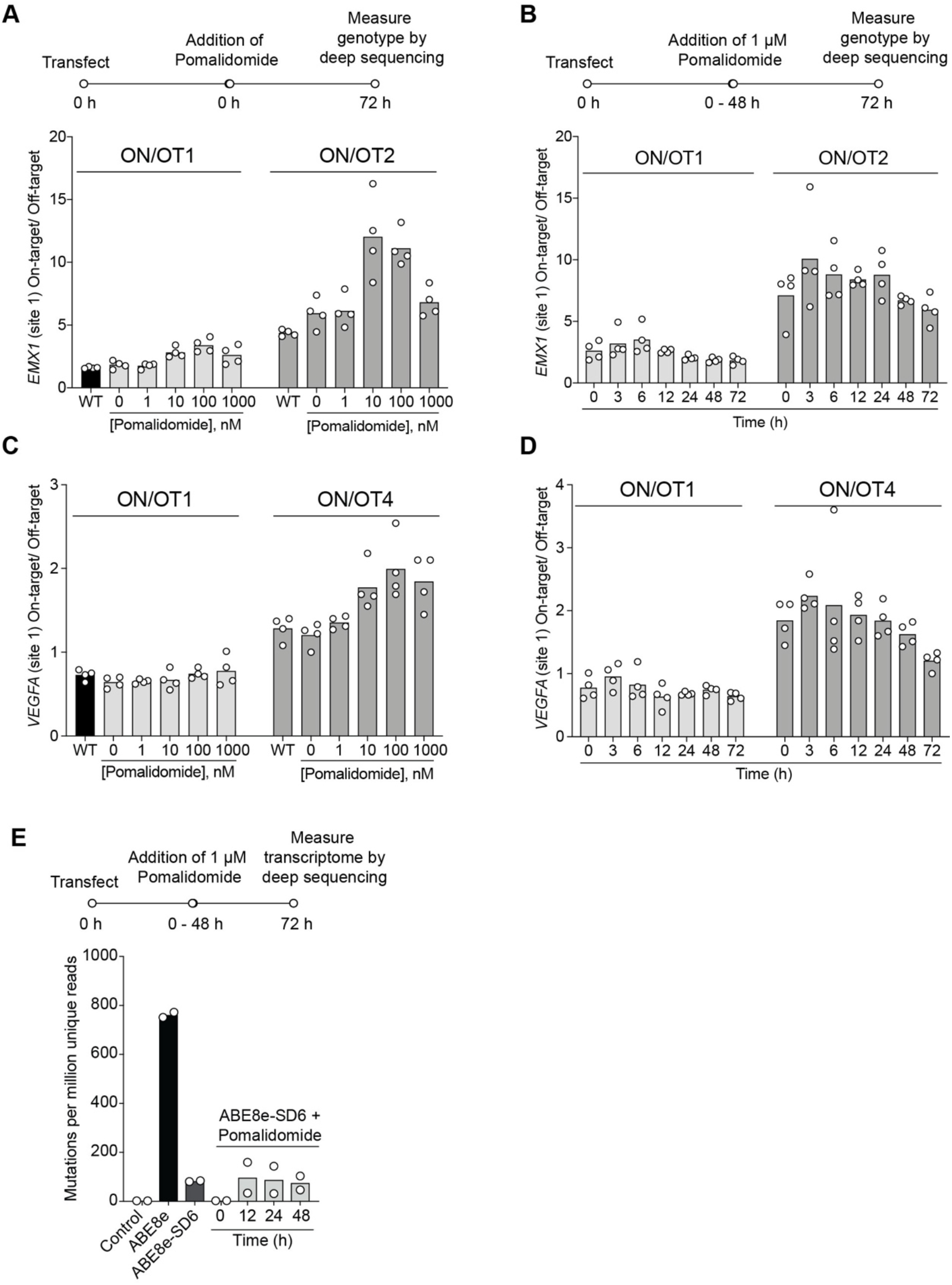
Timely degradation of CRISPR-associated proteins improve the targeting specificity. **(A-D)** Impact of Cas9 half-life on targeting specificity investigated in HEK293T cells. Pomalidomide dose-dependent control (A, C) of on- versus off-target activity of LSD-Cas9 targeting *EMX1* (A), *VEGFA* (C). Pomalidomide-induced half-life-dependent (B, D) control of on- versus off-target activity of LSD-Cas9 targeting *EMX1* (B), *VEGFA* (D). **(E)** Time-dependent control on the transcriptome wide mutations induced by the ABE8e and ABE8e-SD6 upon addition of pomalidomide.

To perform this same test, the on- and off-target specificity of our superdegron base editor system, we transfected HEK293T cells with ABE8e-SD6 and *HBG2*-targeting gRNA and treated the cells with pomalidomide. There was a dose-dependent reduction of Cas9 dependent both on- and off-target editing at the *HBG2* loci, (Figure S5A) and however, no major changes in the targeting specificity for on- versus off-target (Figure S5C). We anticipated that a fast degradation of ABE8e would again allow us to titer the optimal ABE lifetime to maximize the on-target editing, so we tested pomalidomide treatment at different time points. Limiting the base editor activity to short lifetime decreased the editor activity (Figure S5B). However, we did in fact observe a small enhancement of the on- versus off-target ratio with a shortened ABE lifetime (Figure S5D), as limiting the ABE lifetime to within 24 hours showed an increase in target specificity compared to WT ABE. These no or small changes in the Cas9 dependent on-target versus off-target actions may be due to the dosage of editor as well as fast and high deaminase (TadA) activity of ABE8e. To understand the effect of base editor’s lifetime on transcriptome wide mutations, we performed RNAseq analysis. Introduction of the superdegron itself dramatically reduced the mutations in RNA, providing little dynamic range to allow pomalidomide-mediated degradation to have an effect on the transcriptome wide mutations (Figure 5E). However, additions of pomalidomide at lower time point (0 h) abolished the mutations presumably due to low expression of the editor (Figure 5E). Thus, for adenine base editors, the fusion of response element not only allows control of half-life but also dramatically reduces RNA off-target.

### Demonstration of *in vivo* control of base editor’s activity

In a step towards *in vivo* control of base editor activity, we generated an AAV bearing the adenine base editor. Here, we used a split base-editor/dual-AAV strategy, in which the adenine base editor is divided into an amino-terminal (1–573 aa) and carboxy-terminal half (574–1368 aa) with the response element present at loop-231 as described above and each base-editor fragment bearing split intein.^42^ Following co-infection by AAV9 particles expressing each base-editor/split-intein half, protein splicing in trans reconstituted the full-length base editor (Figure 6A).^42^ Because our degradation system requires the human CRBN component, we established the effect of pomalidomide on base editor activity in a human CRBN knock-in C57Bl6/J mice.^43^ We used ABE8e-SD6 AAV9 particles targeting the *Pcsk9* site in the liver.^44,45^ We injected a dose of 5 × 10^11^ AAV9 N-terminal and C-terminal ABE8e-SD6 particles (a 1:1 mixture at 2.5×10^11^ vg each AAV) via retro-orbital eye injection and allowed the base editor to self-assemble and edit the genome for the first week. For the following two weeks, we administered pomalidomide daily at an oral dose of 30 mg/kg. We euthanized the mice at three weeks post viral-particle injection and harvested the liver and blood (Figure 6B). The base editing activity of ABE8e-SD6 was analyzed by next-generation sequencing after extracting the genomic DNA from the liver. Here, the mice administered with pomalidomide showed significantly lower base editing activity compared to the control, validating the degradation of base editors in the mouse livers (Figure 6C). ELISA-based Pcsk9 levels in the plasma of pomalidomide-treated mice were also higher compared to the control mice (Figure 6D), suggesting the usability of our system to control base editor activity *in vivo.*

**Figure 6.**
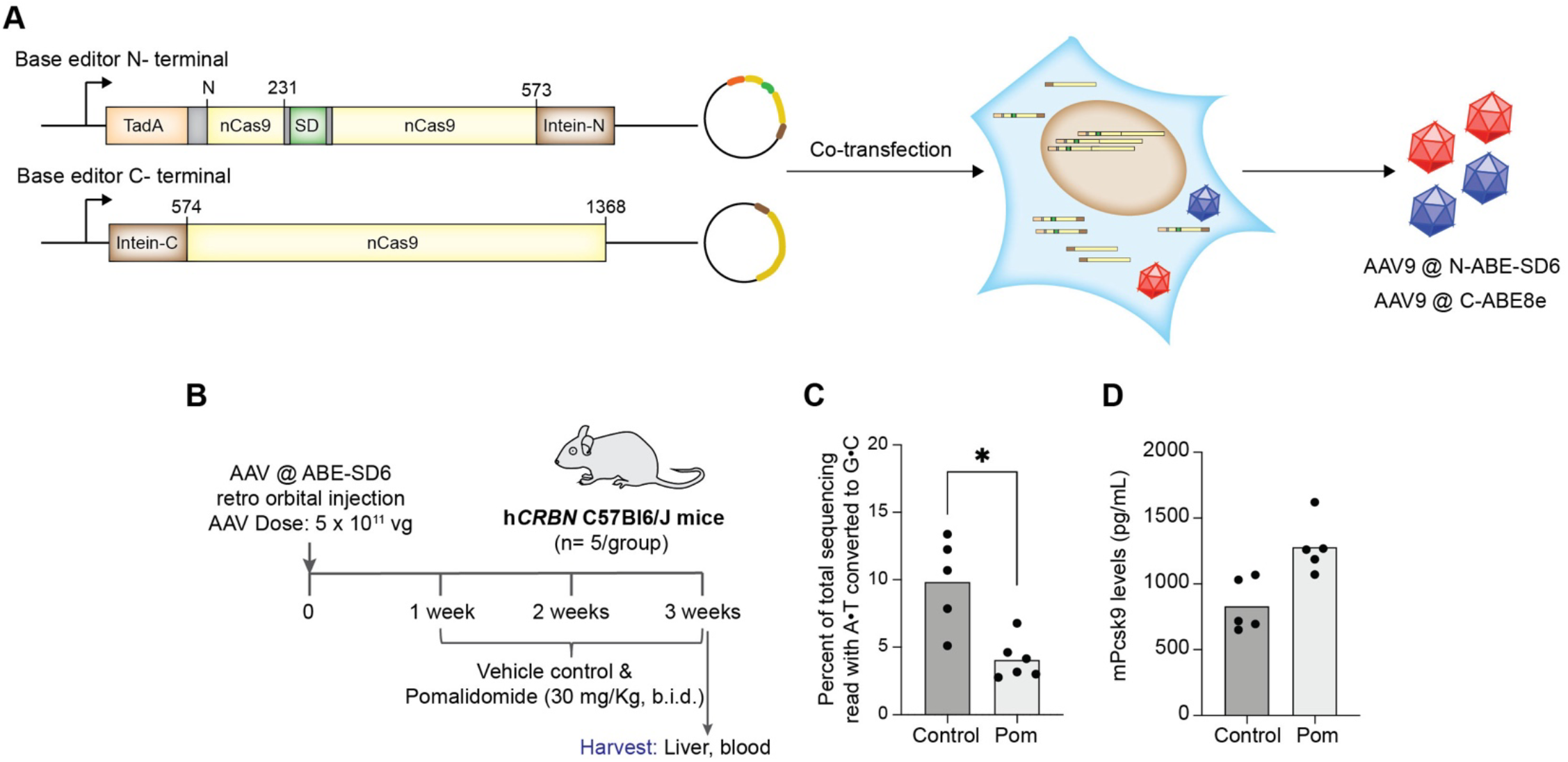
Superdegron applicability and degradation demonstration of genome editing proteins in mouse models. **(A)** Intein reconstitution strategy uses two fragments of protein fused to halves of a split intein that splice to reconstitute a full-length protein following co-expression in host cells. **(B)** Schematic showing retro-orbital injection of 5 × 10^11^ vg of AAVs consisting of split ABE-SD6 in humanized CRBN knock-in C57Bl6/J mice. AAV-injected mice were given one week for genome editing before pomalidomide or the vehicle control were administered orally for two weeks. After three weeks of AAV injection, mice were euthanized, and their liver and blood were harvested to analyze the base-editing levels in the liver and the Pcks9 levels in plasma. **(C, D)** NGS-based analysis showing (C) the conversion of AoT to GoC in the livers and (D) Pcks9 levels in the blood plasma of control and pomalidomide-treated mice.

## Discussion

Of the three elements of central dogma (Proteins, RNA, and DNA), nearly all therapeutic agents target proteins. CRISPR-based technologies provide a novel therapeutic modality by expanding the target scope to genome’s coding and non-coding regions. However, these systems do not display attributes of a typical therapeutic agent,^1, 6^ including precision control of dosage and half-life, and in many cases, the activity of these systems is described as “genome vandalism.” Molecular glues are a rising therapeutic modality used for precision control of dose and half-life of proteins. These glues trigger target protein degradation by bringing together the target protein and ubiquitin ligase, consequently triggering protein ubiquitination and subsequent degradation. In this study, we leverage the ascendant molecular glue technologies to bring the desired attributes of a therapeutic agent—dose, temporal-, and half-life-control—to the realm of CRISPR-based technologies. The small footprint of the response element did not perturb the activity of CRISPR-based technology and allowed facile packaging in contemporary AAV-based delivery systems. Furthermore, the controller is FDA-approved drug pomalidomide with desirable pharmacokinetic and pharmacodynamics properties and the response element, being of human origin, should not be immunogenic. Our system was efficacious in diverse cell types, including stem cells, and easily transportable from Cas9 to CRISPR-based technologies.

We were not only able to switch off and fine-tune the activities of CRISPR-based technologies but also build an on-switch by degrading anti-CRISPR proteins. While several on-switches have been developed for CRISPR-associated nucleases,^13, 28, 45, 46^ similar switches are missing for CRISPR-based technologies, including base editors, perhaps because of size/site limitations for appending regulatory domains on these technologies, which already have multiple fusions at the termini. Constitutively active base editors can lead to unintended genomic alterations and transcriptomic changes, which raises significant biosafety concerns in therapeutic genome editing.^47–49^ We could switch on, fine-tune, and switch-off CRISPR-based technologies using pomalidomide as a controller. This fine-tuning allowed us to precisely control on-target vs. off-target editing ratio, modulate the genotypic outcome, and control base editor activity *in vivo.*

The existence of Cas9-reactive T_effector_ cells and antibodies is a major concern. Our system can allow modulation of the efficacy versus immunogenicity and provide a “kill-switch” in the event of severe adverse reaction (e.g., cytokine storm) as was observed in the first gene therapy clinical trials. Furthermore, fine-tuning the Cas9 halflife would ameliorate the genotoxicity, as constitutively active Cas9 can cause unwanted large deletions and complex genomic rearrangements in edited cells, which could have pathogenic consequences, especially in mitotically active cells.^50–53^ Following Cas9-induced double-strand breaks, the nature of DNA repair pathways determines the gene-editing outcome. Since Cas9 remains bound to the double-strand break and potentially impacts the recruitment of DNA repair machinery, controlling Cas9 half-life will allow control over the nature of the genotypic outcome.^17, 54^ In the absence of such control, the genotypic output from a Cas9-mediated gene knock-out will be highly heterogeneous and may result in the generation of several neo-epitopes, further aggravating the immunogenicity problem. Our system allows precise control of the genotypic outcome by controlling Cas9 half-life. Finally, while we have focused our efforts on Cas9 derived from *S. pyogenes,* several next-generation CRISPR-associated nucleases with superior attributes are emerging and we anticipate that our system will be easily transportable to these emergent nucleases. Thus, we anticipate that this system to precisely control the dosage and half-life of CRISPR-based technologies will propel the development of therapeutic applications of genome editing technologies.

## Experimental methods

### Reagents and plasmids

Lipofectamine 3000 (Invitrogen, Carlsbad, USA) was used as transient transfection agent following the manufacturer’s protocol. TransIT-LT1 transfection reagent (Mirus Bio) and Polybrene Infection/Transfection Reagent (EMD-Millipore) was used for lentiviral transfection. An SF, SE and P3 Cell Line 4D-Nucleofector X Kit (Lonza) was used as the nucleofection agent following the manufacturer’s protocol. All the plasmids are submitted to addgene will be available to researchers upon request, and sequences of different superdegron constructs is included in the Supporting Information.

### Cell Culture

HEK293FT cells, U2OS, and U2OS.eGFP-PEST cells were cultured in Dulbecco’s modified Eagle’s medium (DMEM; Invitrogen), and Jurkat cells were cultured in RPMI-1640 medium (RPMI; Gibco) supplemented with 10% fetal bovine serum (FBS; Gibco), 1x penicillin-streptomycin (Penstrep; Gibco) and 1x sodium pyruvate (Gibco) at 37°C in a 5% CO_2_ atmosphere. The stable iPSCs expressing the CRISPR repressor were cultured in a xeno-free and feeder–cell-free state in Essential 8™ Flex Medium (Gibco) along with the addition of Essential 8™ Flex Supplement (50X) supplied by manufacturer. iPSC cells were subcultured according to the protocol mentioned for Gibco A18945. Cells matched their expected cell-type morphology and were routinely maintained at <90% confluency.

### Generation of stable cell lines

U2OS cells were nucleofected with 50 pmol of Cas9 RNP containing the gRNA-targeting AAVS1 locus along with the AAVS1 donor plasmid^39^ containing N-HiBiT-LSD-Cas9-P2A-eGFP. They were cultured via puromycin selection and sorted for eGFP positive population via FACS (Sony SH800 cell sorter). Jurkat cells were infected with lentiviral particles containing the LSD-Cas9-FLuc-IRES2-*eGEP* transgene, and the expanded eGFP-expressing cells were sorted by Sony SH800 cell sorter. U2OS stable cells expressing N-HiBiT-LSD-Cas9-P2A-eGFP were infected with lentiviral particles containing the LgBiT-IRES2-mCherry transgene, and the expanded cells were sorted by Sony SH800 cellsorter for double-positive eGFP and mCherry expression. HEK293T cells were infected with lentiviral particles carrying ABE8e-SD6-IRES2-eGFP or LSD-dCas9-KRAB-MeCP2 transgenes. Expanded cells were sorted either sorted for the eGFP expression by Sony SH800 cell sorter or blasticidin resistance marker expressing cells. iPSCs were transfected with LSD-dCas9-BFP-KRAB and TALENS targeting the human *CLYBL* safe harbor locus (between exons 2 and 3) (pZT-C13-R1 and pZT-C13-L1, Addgene #62196, #62197) using transfection reagent, DNA In-Stem (VitaScientific). After 14 days, BFP-positive iPSCs were isolated via Sony SH800 cellsorter.^35^ HEK293FT cells were nucleofected with 50 pmol of Cas9 RNP containing the gRNA targeting the AAVS1 locus^39^ and AAVS1 donor plasmid containing 3xFLAG-CSD-AcrIIA4-P2A-eGFP. The cells were cultured via puromycin selection and were sorted for eGFP positivity via FACS (Sony FACS sorter).

### Immunoblotting

U2OS, HEK293T, iPSC, and HEK293FT cells stably expressing LSD-Cas9, ABE8e-SD6, LSD-dCas9-BFP-KRAB, and 3xFLAG-CSD-AcrIIA4 proteins were incubated with different concentrations of pomalidomide for different time points. Cells were pelleted by centrifugation at 1,000 g for 5 min and lysed with M-PER™ mammalian protein extraction reagent (Thermo Scientific) supplemented with Halt™ protease and phosphatase inhibitor cocktail (Thermo Scientific) on ice for 30 min. The total soluble lysate was collected as the supernatant following centrifugation at 20,000 g for 20 min at 4°C. Protein concentration was determined using a bicinchoninic acid assay (BCA assay; ThermoFisher).

Twenty micrograms of total cell lysates were electrophoresed in 7% NuPAGE™ tris-acetate mini protein gel (for Cas9: 150 V for 50 min.) or 4–12% NuPAGE™ bis-tris mini protein gel (for AcrIIA4: 200 V for 50 min.) and transferred onto an iBlot™2 nitrocellulose transfer mini stack membrane (for Cas9) or iBlot™2 PVDF transfer mini stack membrane (for AcrIIA4) using the iBlot2 dry transfer instrument (Thermo fisher scientific). For Cas9 transfer, we used a gradient protocol starting at 20 V for 3 min followed by 23 V for 4 min and then 25 V for 3 min. Similarly, AcrIIA4 membrane transfer was carried out at 20 V for 7 min. Transferred nitrocellulose or PVDF membranes were subjected to immunoblotting with a monoclonal antibody anti-Cas9 antibody (Abcam: ab189380 or ab191468), anti-FLAG antibody (Cell Signaling Technologies: 2368S), and a monoclonal anti-*β*-actin antibody (Cell Signaling Technologies: 3700S or 4970S). Bands were detected with IRDye-labeled infrared secondary antibody (LI-COR Biosciences: IRDye® 680RD donkey anti-rabbit IgG secondary antibody, P/N: 926-68073; and IRDye® 800CW donkey anti-mouse IgG secondary antibody, P/N: 926-32212) on an Odyssey CLx imager (LI-COR Biosciences).

### *eGFP* disruption assay

Approximately 0.25 × 10^6^ U2OS.eGFP-PEST cells were nucleofected with 300 ng of Cas9 expression plasmid and 30 ng of *eGFP*-targeting gRNA expression plasmid using the SE Cell line 4D-Nucleofector X Kit (Lonza) and DN100 pulse program. For anti-CRISPR screening, approximately 0.25 × 10^6^ U2OS.eGFP-PEST cells were nucleofected with 300 ng of Cas9 expression plasmid, 30 ng of *eGFP*-targeting gRNA expression plasmid, and 50 ng of AcrIIA4 expression or 300 ng of Cas9-P2A-CSD-AcrIIIA4 plasmid and 30 ng *eGFP*-targeting gRNA expression plasmid using the SE Cell line 4D-Nucleofector X Kit (Lonza) and DN100 pulse program. Approximately 20,000 transfected cells were seeded per well in 100 μL of DMEM medium in a 96-well plate (Perkin Elmer cell carrier) and incubated for 24 h at 37°C with the indicated amount of pomalidomide. Cells were fixed in 4% paraformaldehyde and stained by HCS NuclearMask Blue stain (Invitrogen). Imaging was performed using an Operetta CLS microscope and the Operetta Harmony 4.8 (PerkinElmer) software was used for the analysis.

### HiBiT-LgBiT complementation-based degradation assay and Immunoblotting

HEK293FT cells (0.5 × 10^6^ cells per well in 12 well plate) were transiently transfected with WT-Cas9 and different Cas9-degron constructs using Lipofectamine 3000. For CRBN mediated degradation studies, HEK293T cells with *CRBN* -/- and *CRBN* +/+ (0.5 × 10^6^ cells per well in 12 well plate) were transiently transfected with N-HiBiT-LSD-Cas9 expressing plasmid using Lipofectamine 3000. Cells were treated with indicated amounts of pomalidomide and incubated for 24 h. After treatment, the cells were washed with DPBS and were pelleted by centrifugation at 1,000 g for 5 min and lysed with M-PER™ mammalian protein extraction reagent (Thermo Scientific) supplemented with the Halt™ protease and phosphatase inhibitor cocktail (Thermo Scientific) on ice for 30 min. The total soluble lysate was collected as the supernatant following centrifugation at 20,000 g for 20 min at 4°C. Protein concentration was determined using a bicinchoninic acid assay (BCA assay; ThermoFisher). For each condition, 10 μg of the protein lysates were diluted to 50 μL using the M-PER buffer followed by complementation with LgBiT using Nano-Glo HiBiT Lytic Detection System (Promega) according to the manufacturer’s protocol. The luminescence signal was recorded by an EnVision Multilabel Plate Reader with EnVision Manager 1.13 (PerkinElmer). Similarly Immunoblotting of N-HiBiT LSD-Cas9 carried out as described earlier in Immunoblotting section.

### NanoBRET based ternary complex formation assay

U2OS cells stably expressing N-HiBiT-LSD-Cas9 (0.8 × 10^6^ cells per well of 6 well plate) were transiently transfected with 100 ng LgBiT plasmid and 2 μg HT-CRBN expressing plasmid using lipofectamine 3000 reagent. Similarly, HEK293T *CRBN* -/- cells (1 ×10^6^ cells per well of 6 well plate) were transiently transfected with 100 ng N-HiBiT-LSD-Cas9, 100 ng LgBiT, and 2 μg HT-CRBN expressing plasmids using lipofectamine 3000 reagent. The following day, 2000 transfected cells were replated into white 384-well tissue culture plates in the presence or absence of 1 μM HaloTag NanoBRET-618 Ligand (Promega) and incubated overnight at 37 °C, 5% CO_2_. The following day, medium was replaced with Opti-MEM (Gibco) containing 20 μM Vivazine, an extended time-released substrate (Promega), and plates were incubated at 37 °C, 5% CO_2_, for 30 min before addition of DMSO or different concentrations of pomalidomide and incubated for 15 minutes. Then the plates were read by an EnVision Multilabel Plate Reader with EnVision Manager 1.13 (PerkinElmer) using two filters, 600nm long pass filter (AT600LP, Ø15 mm, Chroma technologies) and 480/30 nm bandpass filter (2100-5040, barcode 104, Perkin Elmer) for integration time of 1 second. Background subtracted NanoBRET ratios expressed in milliBRET units (mBRET) were calculated from the equation.

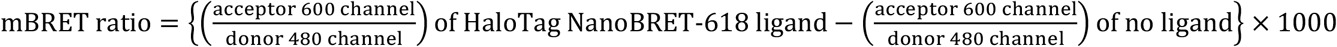

### Screening and next-generation sequencing for base editors ^34^

For screening of the base editor-degron constructs, HEK293T cells (0.5 × 10^6^ cells per well were seeded in a 48-well plate 16h prior to transfection) were transiently transfected with 750 ng of base-editor plasmid and 250 ng of *HBG2*-targeting gRNA plasmid using lipofectamine 2000 in the presence of indicated amount of pomalidomide. Genomic DNA was extracted 72 h after transfection using a crude lysis buffer. Next-generation sequencing samples were prepared via two-step PCR following the protocol reported previously.^53^ Amplicon sequences were analyzed by CRISPResso 2.^54^ For dose-dependent On versus Off target analysis of the ABE-SD6 construct, HEK293T cells (0.5 × 10^6^ cells per well were seeded in a 48-well plate 16h prior to transfection) were transiently transfected with 750 ng of base editor plasmid and 250 ng of *HBG2*-targeting gRNA plasmid using lipofectamine 2000 and were incubated in the presence of pomalidomide. For dose-dependent degradation, different amounts of pomalidomide were added at 0 h after transfection, whereas for time-dependent degradation, 1 μM pomalidomide was added at 0, 3, 6, 12, 24, 48 h post transfection. Genomic DNA was extracted 72 h after transfection using a DNeasy Blood & Tissue Kit (Qiagen), and next-generation sequencing samples were prepared via the two-step PCR protocol that was reported previously.^53^ Amplicon sequences were analyzed by CRISPResso 2.^54^

For AntiCRISPR mediated switch on of base editors, HEK293FT cells (0.5 × 10^6^ cells per well in a 48-well plate 16h prior to transfection) were transiently transfected with 250 ng of ABE8e plasmid, 75 ng of *VEGFA*-targeting gRNA plasmid and 750 ng CSD-AcrIIA4 plasmid or 150 ng CBEmax plasmid, 50 ng of *HEK2* targeting gRNA plasmid along with 250 ng CSD-AcrIIA4 plasmid using lipofectamine 3000 in the presence of indicated amount of pomalidomide. Genomic DNA was extracted 72 h after transfection using a crude lysis buffer. Next-generation sequencing samples were prepared via two-step PCR following the protocol reported previously.^53^ Amplicon sequences were analyzed by CRISPResso 2.^54^

### FACS-based quantification of *TFRC* repression^57^

To quantify the effect of pomalidomide-induced degradation of dCas9 based knockdown efficiency, iPSCs were nucleofected with TFRC-targeting gRNA plasmid and treated with indicating amounts of pomalidomide for 48 h. Approximately 1 x 10^5^ iPSC cells were suspended in PBS, washed once in PBS with 5% FBS, resuspended in 100 μL of PBS containing 5% FBS and blocked with 2.5 μg of Fc-blocking solution (BD Bioscience, Cat #564220) for 15 min at room temperature. Cells were then stained for transferrin receptor expression by adding 0.25 μg of PE/Cy7 anti-human CD71 [CY1G4] (Biolegend, Cat# 334111) directly to the blocking solution for 30 min at room temperature. Excess antibody was removed by two washes in PBS containing 5% FBS, and then cells were re-suspended in 150 μL of PBS with 5% FBS and analyzed on a BD FACS Canto2 flow cytometer (BD Biosciences) for PE-Cy7 expression.

### Gene expression and real-time qPCR studies

HEK293T cells stably expressing LSD-dCas9-KRAB-MeCP2 protein were transfected with 100 ng of gRNA plasmid targeting *CXCR4* or *BRCA1* genes were incubated for 48 h with indicated amounts of pomalidomide. Following 48 h cells were washed with cold PBS and the total RNA was extracted using TaqMan® Gene Expression Cells-to-CT™ Kit as per instruction of the manufacturer. Gene expression was normalized to the housekeeping gene *GAPDH* via the 2^-ΔΔCt^ method.^58^

### U2OS cell culture with reduced-gRNA library ^37^

The development of the U2OS cell line containing the stable Reduced Library genomic integration was described previously.^37^ Briefly, approximately 100,000 U2OS cells were plated per well in a 6-well plate and were selected with 1 μg/mL of puromycin starting 24 h after transfection and continuing for >1 week. Sequential Cas9 plasmid integration was performed using the Lipofectamine 3000 transfection protocol followed by selection with 10 μg/mL of Blasticidin starting 24 hours after a 2-day transient transfection. Next, 1 μM of pomalidomide was introduced at 0, 6, 12, 24, and 48 h timepoints after LSD-Cas9 plasmid transfection. The genomic DNA was collected at 120 hours after transfection.

### Reduced Library and HTS data analysis

The analysis for this paper was performed on 81 M raw reads from Illumina deep sequencing collected following the addition at four time points of the small molecule pomalidomide to a CRISPR library consisting of 48 well-characterized guide sequences. Sequencing reads were filtered to remove low-quality (Illumina average quality < 28) or unmapped reads and were genotyped. For each sequencing read representing a CRISPR-Cas9 cutting event, the cutting genotype was identified and categorized as an insertion or deletion event, and the overall fractions of insertions and deletions of all lengths were computed for two replicates at each time point. The analysis protocol was described previously.^37^

### Droplet digital PCR-based assay^27, 41^

First, 400 ng of LSD-Cas9 expression plasmid along with 40 ng of *RBM20* gRNA plasmid, and 40 pmol of single-stranded oligonucleotide donor DNA (ssODN) were nucleofected to 0.25 × 10^6^ HEK293T cells using the SF Cell Line 4D-Nucleofector kit (Lonza) following the pulse program of CM-130. Transfected cells were plated at 20,000 cells per well in 400 μl of DMEM medium in a 24-well plate (Corning) and were incubated for 72 hours at 37°C with 1 μM of pomalidomide that was added at 0, 3, 6, 12, 24, and 48 h. After 72 hours, the cells were harvested by centrifugation, the genomic DNA was extracted using a DNeasy Blood & Tissue Kit (Qiagen), and the allelic frequency was read by droplet digital PCR.^41^

### HiBiT knock-in assay

HEK293T cells or HEK293T cells with *CRBN+/+* or *CRBN-/-* (approximately 0.25 × 10^6^ cells) were nucleofected with 400 ng of LSD-Cas9 plasmid or 400 ng of Cas9-P2A-CSD-AcrIIA4 plasmid along with 40 ng of *GAPDH-targeting* gRNA plasmid, and 40 pmol of *HiBiT* ssODN using the SF Cell Line 4D-Nucleofector kit (Lonza) following the pulse program of CM-130. Approximately 15,000 cells were seeded in a well in a 96-well plate in 100 μL of DMEM media. Cells were incubated at 37°C for 48 h (for *CRBN-/-* and *CRBN+/+*) or 72 h (HEK293T, time-dependent study) in the presence of the indicated amount of pomalidomide. Next, the cell viability was measured using the PrestoBlue Cell Viability Reagent (Thermo) with SpectraMax M5 and SoftMax Pro 7.0 (Molecular Devices). Next, HiBiT detection was performed using the Nano-Glo HiBiT Lytic Detection System (Promega) according to the manufacturer’s protocol. The luminescence signal was recorded by an EnVision Multilabel Plate Reader with EnVision Manager 1.13 (PerkinElmer). The resulting luminescence signals were normalized with PrestoBlue-based cell viability.

### Next-generation sequencing for Cas9

HEK293T cells (0.15 × 10^6^ cells per well in a 24-well plate) were transiently transfected with 750 ng of Cas9 plasmid and 250 ng of *EMX1(site1*) or *VEGFA (site1*) gRNA plasmid in the presence of pomalidomide. In the dose dependent Cas9 degradation assay, indicated doses of pomalidomide were added from 0 h and incubated until 72 h. In the time dependent Cas9 degradation assay, 1 μM of pomalidomide was introduced at indicated time points after transfection and incubated until 72 h. Genomic DNA was extracted 72 h after transfection using a DNeasy Blood & Tissue Kit (Qiagen). Next-generation sequencing samples were prepared via two-step PCR following the protocol reported previously.^55^ Amplicon sequences were analyzed by CRISPResso 2.^56^

### Next-generation sequencing for Cas9-P2A-CSDAcrIIA4 fusion construct

HEK293T cells (0.15 × 10^6^ cells per well in a 24-well plate) were transiently transfected with 750 ng of Cas9 plasmid, 250 ng of *EMX1(site1)-* targeting gRNA plasmid, and 250 ng of AcrIIA4 plasmid or 750 ng of Cas9-P2A-CSDAcrIIA4 fusion plasmid in the presence of the indicated amounts of pomalidomide. Genomic DNA was extracted 72 h after transfection using a DNeasy Blood & Tissue Kit (Qiagen). Next-generation sequencing samples were prepared via the two-step PCR protocol that was reported previously.^55^ Amplicon sequences were analyzed by CRISPResso 2.

### AAV production^42^

HEK293T/17 cells were maintained in DMEM with 10% FBS without antibiotic in 150-mm dishes (Thermo Fisher Scientific) and passaged every 2–3 days. Cells for production were split 1:3 1 day before polyethylenimine transfection. Then, 5.7 μg of AAV genome, 11.4 of μg pHelper (Clontech) and 22.8 of μg repcap plasmid were transfected per plate. One day after transfection, the media was exchanged for DMEM with 5% FBS. Three days after transfection, cells were scraped with a rubber cell scraper (Corning), pelleted by centrifugation for 10 min at 2,000 g, were resuspended in 500 μL of hypertonic lysis buffer per plate (40 mM Tris base, 500 mM NaCl, 2 mM MgCl_2_, and 100 U/ml salt active nuclease (ArcticZymes), and were incubated at 37°C for 1 h to lyse the cells. The media was decanted, combined with a 5× solution of 40% poly(ethylene glycol) (PEG) in 2.5 M NaCl (final concentration: 8% PEG, 500 mM NaCl), incubated on ice for 2 h to facilitate PEG precipitation, and centrifuged at 3,200g for 30 min. The supernatant was discarded, and the pellet was resuspended in 500 μl of lysis buffer per plate and added to the cell lysate. Incubation at 37°C was continued for 30 min. Crude lysates were either incubated at 4°C overnight or directly used for ultracentrifugation. Cell lysates were gently clarified by centrifugation at 2,000g for 10 min and added to Beckman Quick-Seal tubes via 16-gauge 5” disposable needles (Air-Tite N165). A discontinuous iodixanol gradient was formed by sequentially floating layers: 9 ml of 15% iodixanol in 500 mM NaCl and 1× PBS-MK (1× PBS plus 1 mM MgCl_2_ and 2.5 mM KCl), 6 ml 25% iodixanol in 1× PBS-MK, and 5 ml each of 40 and 60% iodixanol in 1× PBS-MK. Phenol red at a final concentration of 1 μg/ml was added to the 15, 25, and 60% layers to facilitate identification. Ultracentrifugation was performed using a Ti 70 rotor in a Sorvall WX+ series ultracentrifuge (Thermo Fisher Scientific) at 58,600 rpm for 2.25 h at 18°C. Following ultracentrifugation, 3 ml of solution was withdrawn from the 40–60% iodixanol interface via an 18-gauge needle, dialyzed with PBS containing 0.001% F-68, and ultrafiltered via 100-kD MWCO columns (EMD Millipore). The concentrated viral solution was sterile filtered using a 0.22-μm filter, quantified via qPCR (AAVpro Titration Kit version 2; Clontech), and stored at 4°C until use.

### *In vivo* control of base editing in CRBN mouse model^42, 43^

All experiments in live animals were approved by the Dana Farber Cancer Institute institutional animal care and use committee. The Crbn^tm1.1Ble^ C57Bl6/J mouse with Crbn^I391V^ variant was used for the base-editor experiments. Five mice were randomized to each group, and the AAV ABE-SD6 was diluted to 200 μL in 0.9% NaCl for injection. Anesthesia was induced with 4% isoflurane. Following induction, as measured by unresponsiveness to a toe pinch, the right eye was protruded by gentle pressure on the skin, and a tuberculin syringe was advanced with the bevel facing away from the eye into the retrobulbar sinus where N-terminal and C-terminal AAV mix (5 × 10^11^ vg) was slowly injected. Post injection, mice were left for one week, after which we administered 30 mg/kg of pomalidomide orally every day for the next two weeks. After three weeks of AAV administration, the blood was withdrawn by cardiac puncture, and the mice were euthanized to harvest the liver. Pcsk9 from the plasma samples were analyzed by ELISA (R&D systems) as per the manufacturer’s instructions. Genomic DNA from the liver samples was extracted by Beckman Coulter’s genomic DNA extraction kit as per the manufacturer’s instructions. Next-generation sequencing samples were prepared via two-step PCR following the protocol reported previously.^55^ Amplicon sequences were analyzed by CRISPResso 2.^56^

## Acknowledgements

This work was supported by the DARPA (N66001-17-2-4055 to A.C.), NIH (R01GM132825 to A.C.; UG3AI150551, U01AI142756, R35GM118062, and RM1HG009490 to D.R.L.; and R35 HG010717 to L.P.) and HHMI.

## Notes

The authors declare the following competing financial interest: Broad Institute has filed patents claiming inventions to genome editing methods in this manuscript. M.J. has received consulting fees from RA Ventures. M.K. holds equity in and serves on the scientific advisory boards of Engine Biosciences, Casma Therapeutics, Cajal Neuroscience, and Alector and advises Modulo Bio and Recursion Therapeutics. M.K. is an inventor on US patent 11,354,933 related to CRISPRi and CRISPRa screening. L.P. has financial interests in Edilytics, Excelsior Genomics and SeQure Dx. L.P.’s interests were reviewed and are managed by Massachusetts General Hospital and Partners HealthCare in accordance with their conflict-of-interest policies. D.R.L. is a consultant for Prime Medicine, Beam Therapeutics, Pairwise Plants, Chroma Medicine, and Nvelop Therapeutics, companies that use or deliver genome editing or epigenome engineering agents and owns equity in these companies. B.L.E. has received research funding from Celgene, Deerfield, Novartis, and Calico and consulting fees from GRAIL. B.L.E. is a member of the scientific advisory board and shareholder for Neomorph Inc., TenSixteen Bio, Skyhawk Therapeutics, and Exo Therapeutics. A.C. is the scientific founder and scientific advisory board member of Photys Therapeutics.

## Notes

### Summary of Updates

We updated the competing financial interest statement.

