## Supplementary information for "A molecular glue approach to control the half-life of CRISPR-based technologies"

### Figure S1.

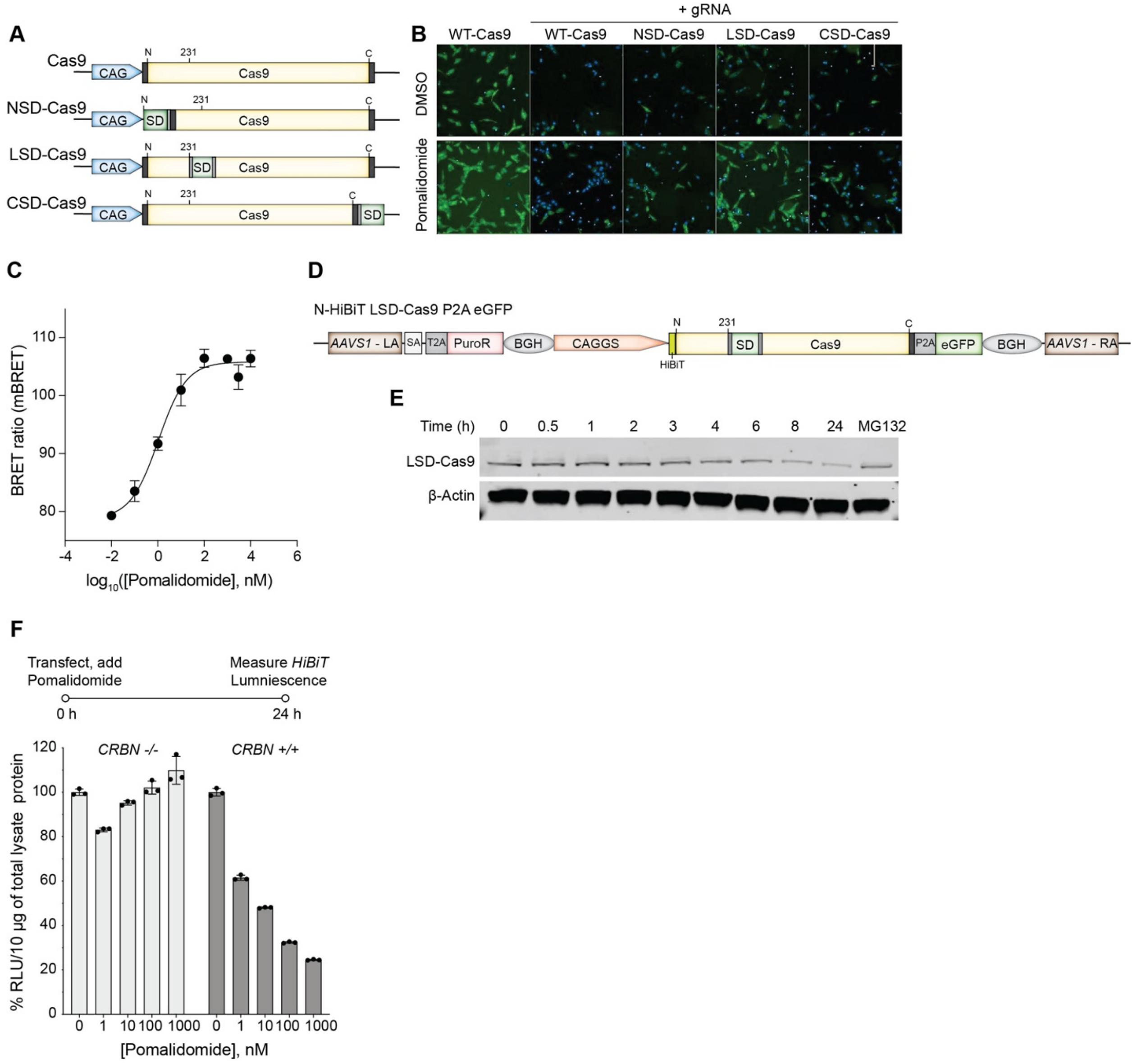


**Figure S1. (A)** Representation of the Cas9 fusions containing a single superdegron tag at the N-terminal (NSD-Cas9), loop-231 (LSD-Cas9), and C-terminal (CSD-Cas9) regions. **(B)** Representative images showing pomalidomide induced degradation of superdegron-Cas9 constructs in U2OS.eGFP.PEST cells in the eGFP disruption assay. **(C)** HEK293T *CRBN*-/- cells were transfected with 1:1:10 ratio of N-HiBiT-LSD-Cas9, LgBiT, and HT-CRBN plasmids for 24 h and followed by incubated with Halotag-618 ligand upon reseeding in 384 well plate. After overnight incubation with Halotag-618 ligand pomalidomide dose were added and incubated for 30 minutes to measure the nanoBRET signal using 480 nm band pass and 615 nm long pass filters on a plate reader (C). milliBRET ratio for the pomalidomide dose-induced ternary complex formation between N-HiBiT-LSD-Cas9, LgBiT, and HT-CRBN in *CRBN*-/- HEK293T cells (D). **(D)** Representation of DNA donor construct usedto knock-in the N-HiBiT-LSD-Cas9 in *AAVS1* locus using Cas9 RNP method. **(E)** Immunoblots showing the pomalidomide-induced time dependent degradation of LSD-Cas9 in transiently transfected HEK293T cell line. **(F)** Pomalidomide-induced degradation of N–HiBiT-fused LSD-Cas9 in transiently transfected HEK293T *CRBN*−/– and *CRBN*+/+ cell lines, measured by HiBiT luminescence.

**Figure S2.**

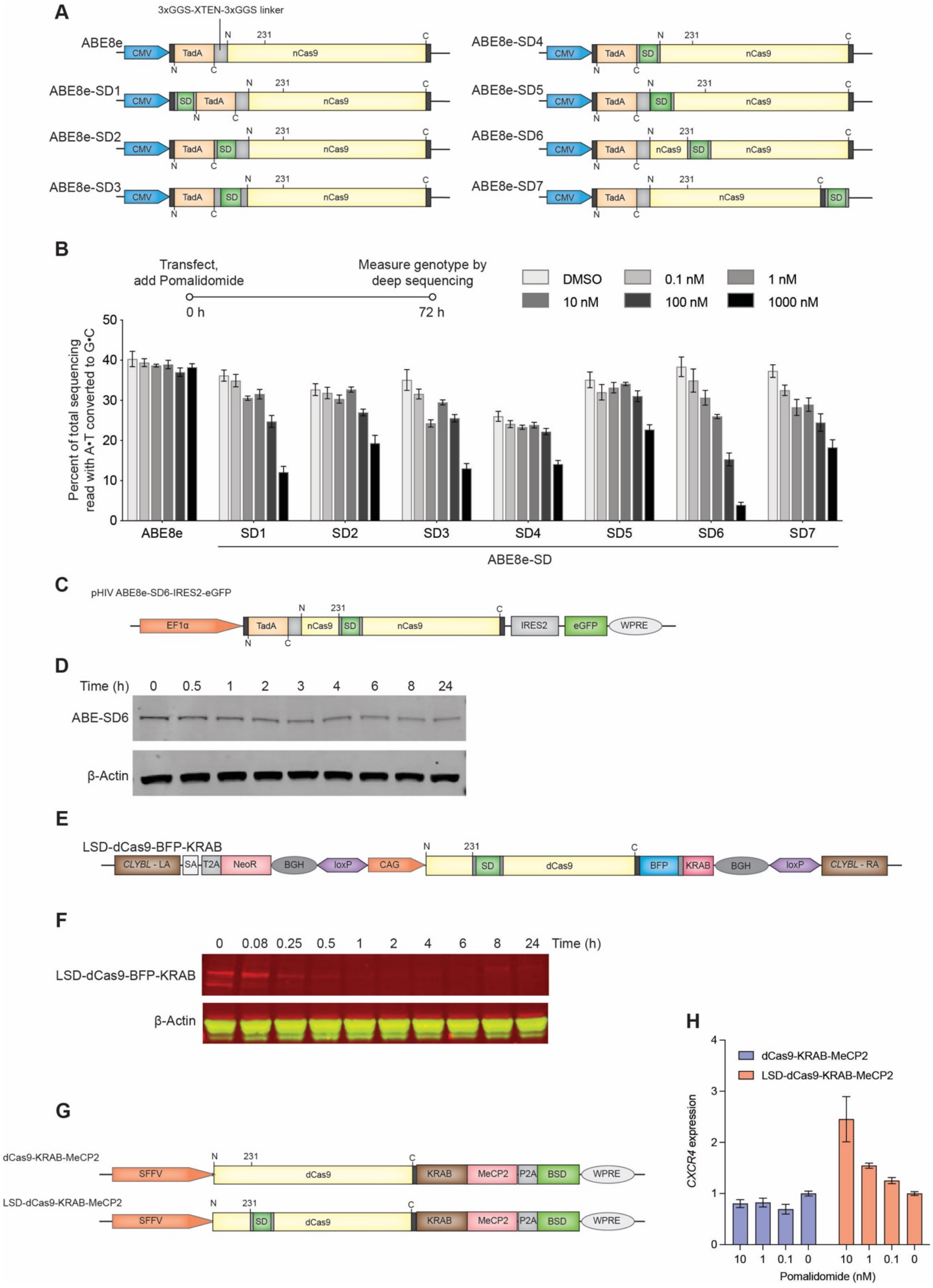


**Figure S2. (A)** Schematic of constructs of adenine base editor (ABE8e) fused with a single superdegron tag at the N-terminus (ABE8e-SD1) or C-terminus (ABE8e-SD2) of TadA deaminase; at the linker region (ABE8e-SD3, ABE8e-SD4); and at the N-terminus (ABE8e-SD5), loop-231 (ABE8e-SD6), and C-terminus (ABE8e-SD7) of the Cas9 nickase. **(B)** Pomalidomide dose-induced base-editor degradation in HEK293T cells transiently transfected with WT ABE8e and ABE8e-superdegron constructs. After 72 h of transfection and pomalidomide treatment, genomic DNA extracted was analyzed by NGS for the conversion of A•T to G•C. **(C)** Schematic of constructs of adenine base editor (ABE8e) fused with a single superdegron tag at loop-231 (ABE8e-SD6), used to make the lentiviral mediated stable cell line generation in HEK293T cells. **(D)** Pomalidomide time-dependent ABE8e-SD6 degradation in transiently transfected HEK293T cells was monitored by immunoblots. **(E)** Schematic of LSD-dCas9-BFP-KRAB system used to knock-in the LSD-dCas9-BFP-KRAB in iPSCs using TALEN method. **(F)** Pomalidomide time-dependent LSD-dCas9-BFP-KRAB- degradation in stably expressing dCas9 in iPSCs was monitored by immunoblots. **(G)** Schematic of dCas9-KRAB-MeCP2 and LSD-dCas9-KRAB-MeCP2 system used to make the lentiviral mediated stable cell generation in HEK293T cells. **(H)** A functional analysis of stably expressing repressor constructs in HEK293T cells upon pomalidomide-dependent degradation was carried out in HEK293T cells by gene expression of *CXCR4* measured by qRT-PCR.

**Figure S3.**

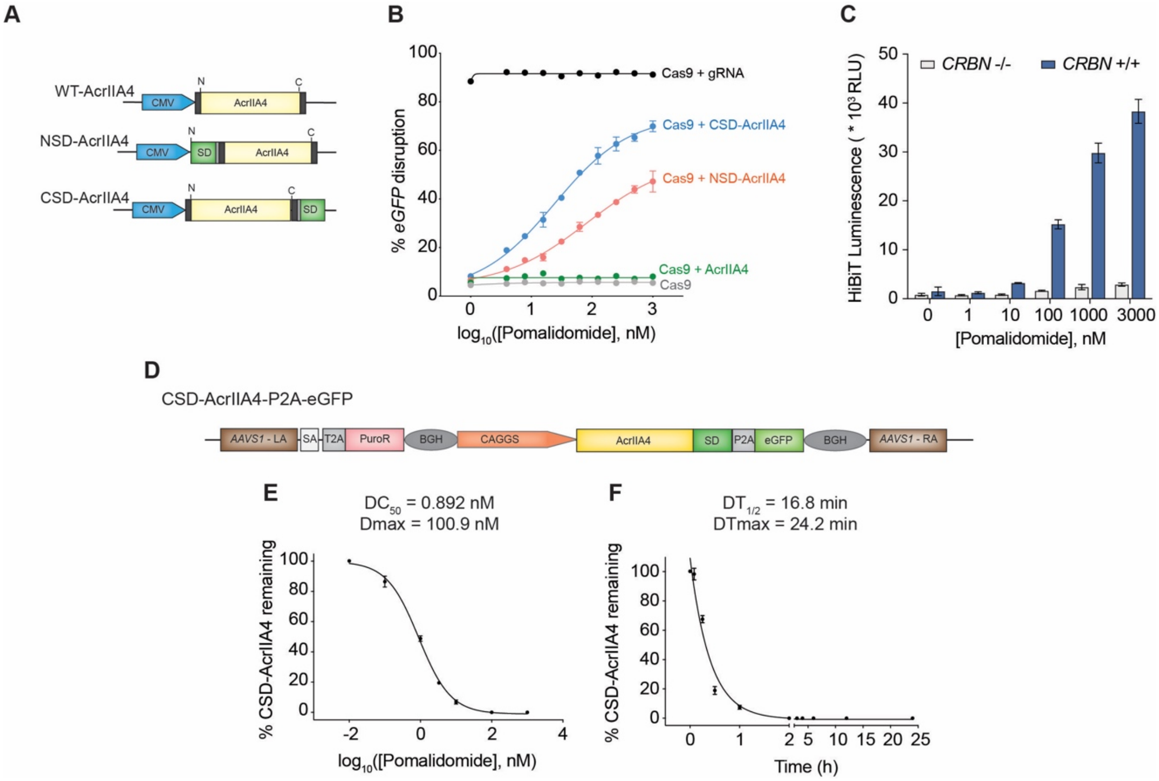


**Figure S3. (A)** Schematic of AcrIIA4 fused with the superdegron. AcrIIA4 is fused with a single superdegron tag at the N-terminal (NSD-AcrIIA4) and C-terminal (CSD-AcrIIA4) regions. **(B)** The fusions were investigated for pomalidomide-induced degradation using the *eGFP*-disruption assay. **(C)** *HiBiT* knock-in in *GAPDH* locus using Cas9-P2A-CSD-AcrIIA4 construct: Pomalidomide dose-dependent (0, 1, 10, 100, 1000, 3000 nM) increase in luminescence activity of HiBiT-LgBiT in *CRBN* +/+ cells but no change in the luminescence levels in the *CRBN* -/- cells, indicating that CRBN mediates the degradation of AcrIIA4 that activates the Cas9 for knock-in of *HiBiT*. **(D)** Schematic of construct used to make HEK293FT cells stably expressing CSD-AcrIIA4-P2A-eGFP from AAVS1 locus investigated for pomalidomide-induced degradation. **(E, F)** Densitometric quantification of pomalidomide-induced dose-dependent (E) and time-dependent (F) degradation of CSD-AcrIIIA4.

**Figure S4.**

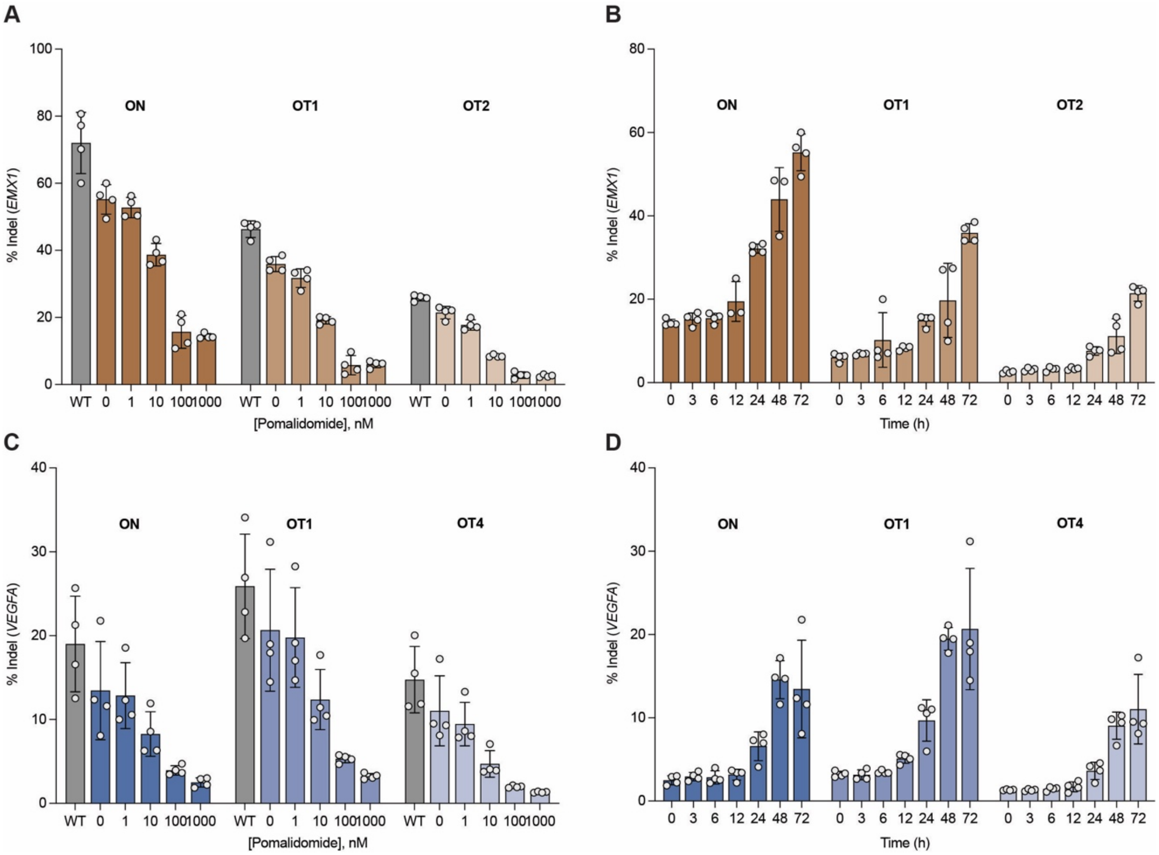


**Figure S4. (A-D)** Impact of Cas9 lifetime on indel formation was investigated in HEK293T cells. Pomalidomide dose-dependent control (A, C) of indel formation by LSD-Cas9 targeting *EMX1* (A), *VEGFA* (C). Pomalidomide-induced lifetime-dependent (B, D) control of indel formation by LSD-Cas9 targeting *EMX1* (B), *VEGFA* (D).

**Figure S5.**

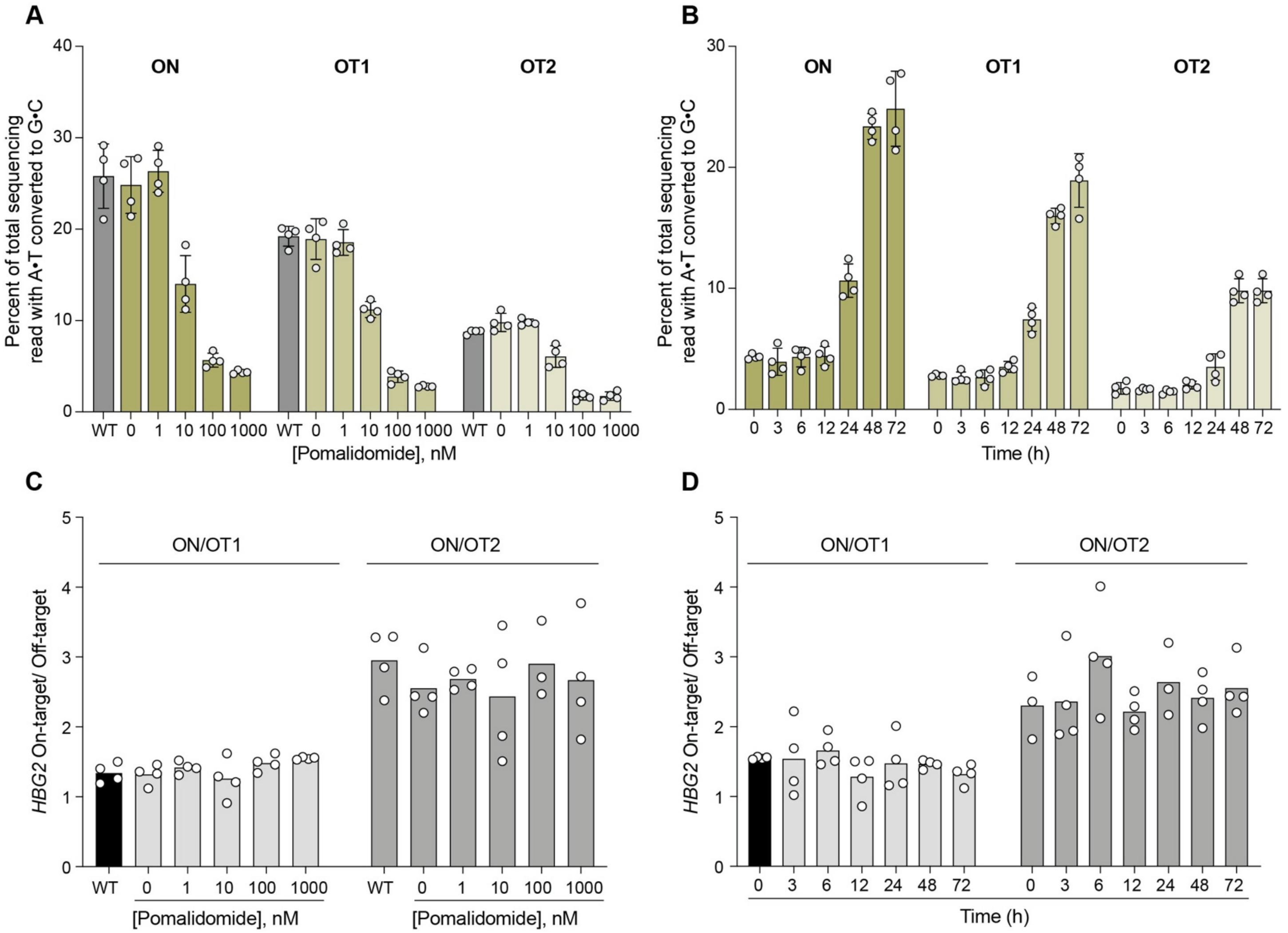


**Figure S5. (A-D)** Base editor lifetime can impact editing specificity. Pomalidomide dose-dependent control (A) and time dependent control (B) on A.T to G.C conversion by ABE8e-SD6 targeting *HBG2* gene. On-target and two off-target sequences were measured. Pomalidomide dose-dependent control (C) and time dependent control (D) of on- versus off-target activity of ABE8e-SD6 targeting *HBG2* gene.

**Figure S6.**

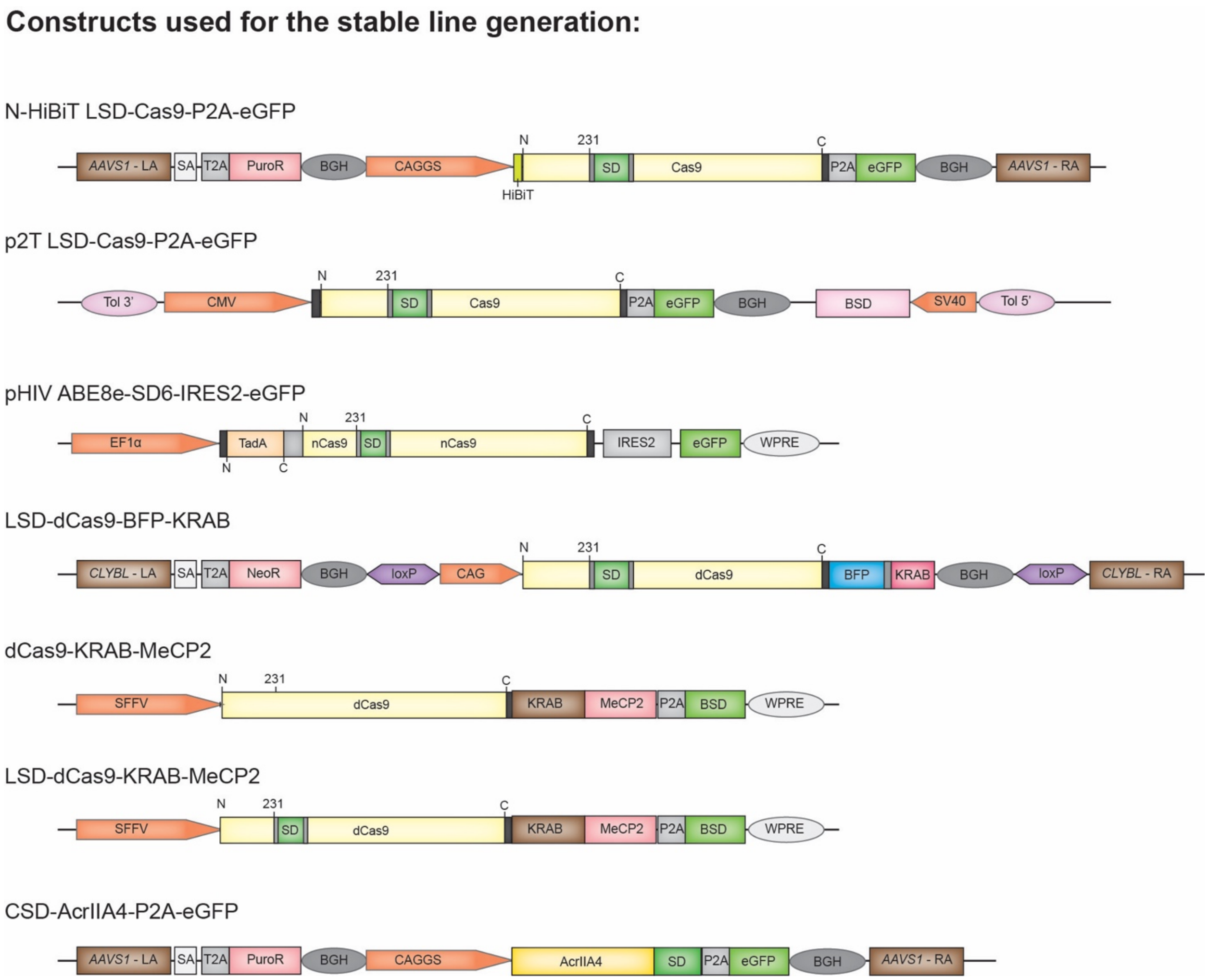


**Figure S6. Constructs used for the generation of stable cells.**
